# Fold-change detection of NF-κB at target genes with different transcript outputs

**DOI:** 10.1101/339697

**Authors:** V. C. Wong, R. Ramji, S. Gaudet, K. Miller-Jensen

## Abstract

The transcription factor NF-κB promotes inflammatory and stress-responsive gene transcription across a range of cell types in response to the cytokine tumor necrosis factor-α (TNF). Although NF-κB signaling exhibits significant variability across single cells, some target genes exhibit fold-change detection of NF-κB, which may buffer against stochastic variation in signaling molecules. However, this observation was made at target genes supporting high levels of TNF-inducible transcription. It is unknown if fold-change detection is maintained at NF-κB target genes with low levels of TNF-inducible transcription, for which stochastic promoter events may be more pronounced. Here we used a microfluidic cell-trapping device to measure how TNF-induced activation of NF-κB controls transcription in single Jurkat T cells at the promoters of integrated *HIV* and the endogenous cytokine gene *IL6*, which produce only a few transcripts per cell. We tracked TNF-stimulated NF-κB RelA nuclear translocation by live-cell imaging and then quantified transcript number by RNA FISH in the same cell. We found that TNF-induced transcription correlates with fold change in nuclear NF-κB with similar strength at low versus high abundance target genes. A computational model of TNF-NF-κB signaling, which implements fold-change detection from competition for binding to κB motifs, was sufficient to reproduce fold-change detection across the experimentally measured range of transcript outputs. Nevertheless, we found that gene-specific trends in transcriptional noise and levels of promoter-bound NF-κB predicted by the model were inconsistent with our experimental observations at low abundance gene targets. Our results reveal a gap in our understanding of RelA-mediated transcription for low abundance transcripts and suggest that cells use additional biological mechanisms to maintain robustness of NF-κB fold-change detection while tuning transcriptional output.

## Introduction

Tumor necrosis factor alpha (TNF) acts on a wide range of cell types and is a critical mediator of inflammatory and stress responses (1) and of the development of certain immune cells, including T lymphocytes (2, 3). TNF stimulation activates a signaling cascade that disrupts the interaction between the transcription factor nuclear factor-κB (NF-κB) and the inhibitor of nuclear factor-κB alpha (IκBα), thereby releasing NF-κB and allowing it to accumulate in the nucleus. NF-κB can then regulate the expression of hundreds of genes, including its own negative regulators *NFKBIA* (encoding IκBα) and *TNFAIP3*, and inflammatory cytokines *TNF* and *IL6* (4). Given its central role in development and immunity, understanding how cells decode responses to TNF signals is of fundamental interest.

Dynamic signaling and response measurements from single cells provide direct information about input-output signal-response functions (5, 6). By measuring dynamic nuclear translocation of the NF-κB RelA subunit and transcription in the same single cells, we previously demonstrated that maximum fold change in nuclear RelA predicts transcript abundance of *NFKBIA, TNFAIP3*, and *IL8* following TNF treatment in HeLa cells, a cell line of epithelial origin (7). This observation had significant implications for our understanding of RelA-mediated signaling in these cells because only two signaling circuits commonly observed in biological systems allow for fold-change detection (8). One is an incoherent type 1 feed-forward loop (I1-FFL), in which a molecular species increases with the input and also inhibits the output (9). By adapting a mathematical model of TNF-stimulated NF-κB RelA signaling (10), we showed that we could accurately reproduce fold-change detection for the measured RelA-target genes by incorporating an I1-FFL mechanism via competition for binding to κB motifs (7). However, the generality of this mechanism across cell types and promoters has not been explored.

NF-κB is also a positive transcriptional regulator of human immunodeficiency virus-1 (HIV), which infects CD4+ T cells (11). We recently observed that TNF-activated NF-κB deterministically controls activation of an HIV reporter, even for latent viruses with very low TNF-inducible transcription (12). Interestingly, we observed significant differences in TNF-induced recruitment of RNA polymerase II at the promoters of these low abundance HIV target genes in Jurkat cells as compared to the high abundance target *NFKBIA* (12). Therefore, we were interested to see if the transcription of low abundance target genes also exhibited fold-change detection, and if the I1-FFL model of NF-κB-mediated transcription could accurately describe the HIV transcription patterns we observed in T cells.

These questions are challenging to address in T cells with a single-cell approach because T cells do not adhere to surfaces and are therefore difficult to track when perturbed. Consequently, we developed a workflow whereby we immobilized Jurkat T cells within a microfluidic cell trap array (13), and then tracked NF-κB RelA nuclear translocation by live-cell imaging and quantified transcript number by fluorescence in situ hybridization of single mRNA (smFISH) in the same cell. Using this workflow, we measured whether TNF-induced activation of NF-κB controls transcription in single Jurkat T cells across diverse NF-κB-target promoters, including HIV and endogenous regulatory and cytokine genes, supporting a range of TNF-inducible transcript levels.

We found that all measured NF-κB-target promoters responded to TNF-stimulated fold changes in nuclear RelA even for transcripts induced at an average mRNA level of 6 or fewer transcripts per cell following 2 hrs of TNF treatment. Our computational model of RelA signaling, which incorporates an I1-FFL mechanism via competition for binding to κB motifs, accurately recapitulated fold-change detection at low abundance genes. However, varying transcriptional output by modulating competition strength in the model failed to recapitulate experimentally observed noise trends at low abundance genes. Furthermore, although our model suggests that reduced RelA binding could result in low transcript abundance, our experiments show that RelA binding varies little across the target promoters that we studied. We conclude that cells must use additional biological mechanisms to maintain robustness of NF-κB fold-change detection while tuning transcriptional output.

## Materials and Methods

### Cell culture and pharmacological treatments

Jurkat T cell clones J65c 4.4 and 6.6 were created as previously described (12). Each cell line contains the same retroviral integration of a full-length human RelA fused at its N-terminus to mCherry (Ch-RelA) and a unique integration of a full-length, non-replication competent reporter virus construct (psLTR-Tat-GFP) in which Nef has been replaced with GFP and the open reading frames of all other viral proteins except Tat are disrupted with premature stop codons. J65c cells were cultured in Roswell Park Memorial Institute 1640 (RPMI) medium (Thermo Fisher Scientific). All media was supplemented with 10% fetal bovine serum (Atlanta Biologicals), 100 U/mL penicillin and 100 μg/mL streptomycin (Thermo Fisher Scientific). Cells were maintained in 5% CO_2_ at 37°C and were never cultured beyond passage 12. J65c cells were grown to 5×10^5^ cells/mL before treatment with recombinant human TNF. As in a previous study, we used 160 ng/ml cycloheximide (CHX; Sigma) to block HIV Tat protein production and the resulting amplification of HIV gene expression (12) and allow us to restrict our analysis to RelA-mediated transcription. This level of CHX is very low and does not change TNF-stimulated RelA translocation dynamics (12) and for consistency, we used these same conditions throughout this study, irrespective of which transcript was assayed.

### Microfluidic device fabrication

The silicon master was etched as previously described (13). Following silanization (see (13) for conditions), a mixture of PDMS base and curing agent (Sylgard 184 Elastomer kit, Dow Corning) was mixed 10:1 by weight and poured over the master. This was degassed and cured for 2 hours at 75 °C. The cured PDMS mold was peeled from the master and cut to fit a #1 glass coverslip. A 7 mm Harris Uni-Core punch (Ted Pella Inc.) was used to punch holes for reservoirs at the inlet and outlet ports. The mold and a #1 25 mm × 75 mm microscope coverslip were rinsed with isopropanol, and dried and cleared of dust with filtered air before being placed inside a PE-25 benchtop plasma cleaner (Plasma Etch). The coverslip and PDMS mold were exposed to O2 plasma at 150 W, 200 mTorr for 2 minutes and adhered immediately upon removal from the plasma chamber. The channel and the inlet and outlet reservoirs were filled with deionized water to maintain hydrophilic surfaces.

### Cell loading protocol

For each experiment, 5×10^6^ cells were suspended in 1 mL of fresh RPMI media prior to being loaded into the microfluidic device. Cell clumps were dispersed immediately before loading by pipetting the cell suspension vigorously and filtering cells through a 35 mm nylon mesh strainer cap tube (BD Falcon). 50 μL of cells were loaded into the inlet and allowed to flow through the channel for 10 min. Flow-through in the outlet was first removed to prevent backflow that can dislodge cells. The cell suspension was then removed from the inlet by gently scraping the bottom of the inlet reservoir with a pipette while withdrawing the media. 100 μL of media was added to the inlet to wash the cells for 3 min to remove any free-floating cells from the main channel, and inlet and outlet reservoirs; this wash step was repeated three times. 50 μL media was added to the inlet after final wash and device was moved onto microscope stage for imaging and stimulation (see below). The continuous flow was maintained throughout the imaging to prevent cells from dislodging from the traps.

### Live-cell imaging

Cells were imaged on a Nikon Eclipse Ti spinning disk confocal microscope (Yokogawa CSU-W1 spinning disk) with a plan apochromatic 60x oil objective (Nikon; NA 1.4). Live-cell imaging was performed on an environmentally controlled stage at 37°c and 5% CO2 (Tokai Hit). Once the loaded device was secured to the stage insert, five regions of interest were identified and their positions relative to fiducial markers on the outside of the channel were recorded. 12 μm Z-stacks with 1 μm slices were recorded for each position as a reference image. Fluid was removed from the outlet and then inlet and 50 μL of RPMI containing TNF was added to the inlet and imaging immediately started. Based on our previous study, we estimated that TNF flows through the entire length of the channel within 30 s and that 50 μL of media was sufficient to maintain a nearly constant flow velocity of 300 μm/s for 2 hours (13). Z-stacks of each region of interest were recorded every 3 min for 110 min after stimulation. After 110 minutes of imaging, cells were fixed by removing fluid from the outlet and then inlet and replacing with 50 μL of 4% formaldehyde in phosphate buffered saline (PBS) for fixation prior to *in situ* hybridization.

### Single molecule RNA fluorescence in situ hybridization (smFISH)

Stellaris RNA FISH probe sets (Biosearch Technologies) targeting *NFKBIA*/IκBα, *TNFAIP3*/A20, *env-GFP*/HIV, and *GAS6* were previously described (7, 12) (14). The *IL6* probe set was designed using online software (biosearchtech.com) and previously described strategies (7, 15). The final probe set for IL6 consisted of 39 probes (Table, below).

**Table.**
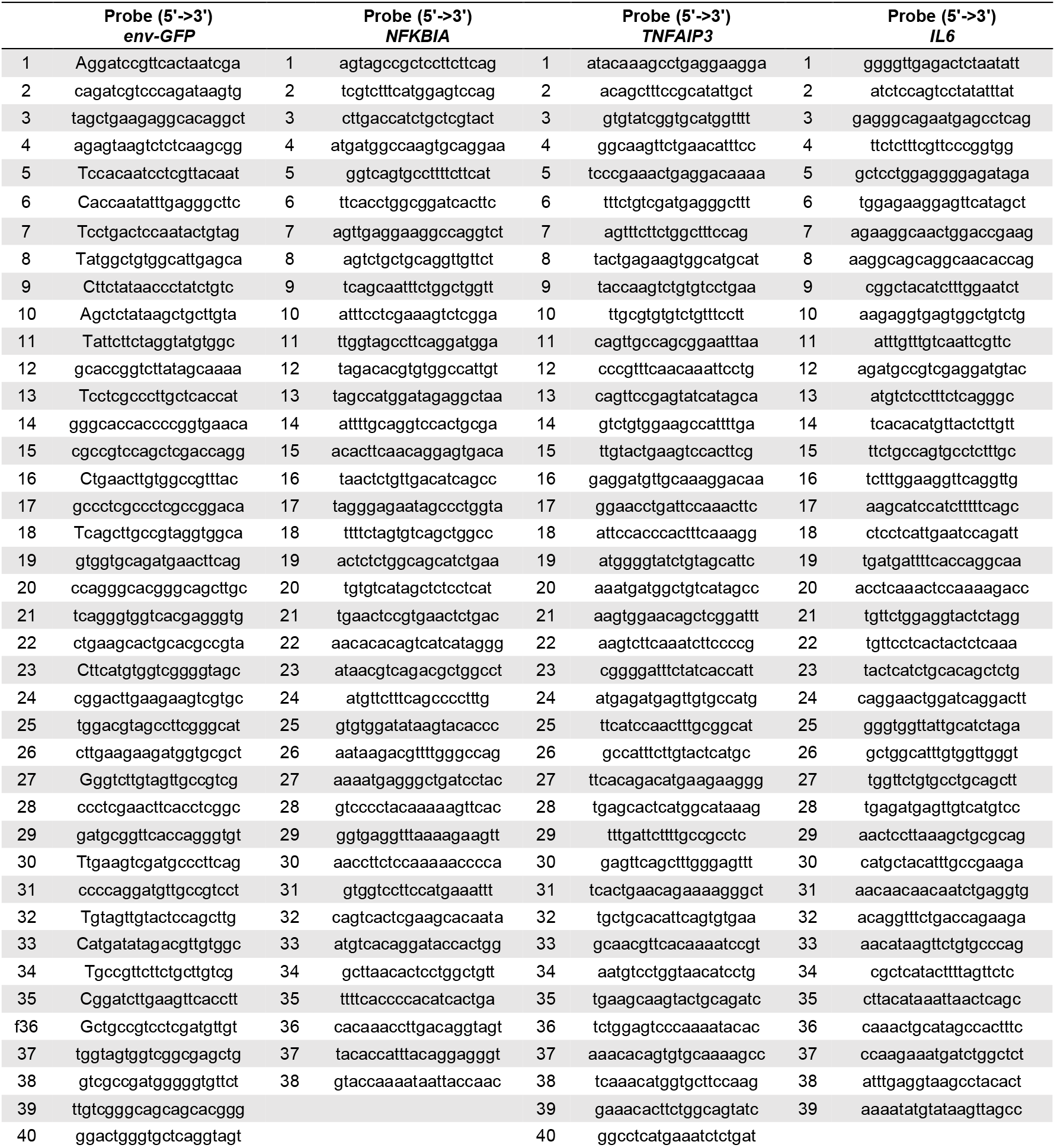
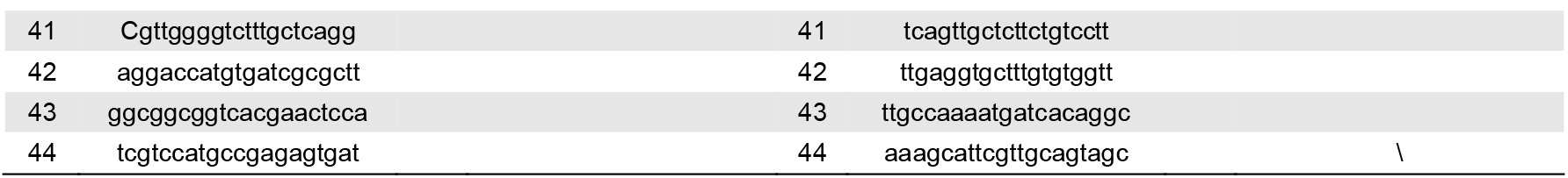
smFISH probes: *env-GFP, NFKBIA*, and *IL6* probes are conjugated to FAM; *TNFAIP3* is conjugated to Quasar 670.

After fixation in the passive-flow device, cells were washed 3 times with PBS, allowing the buffer to flow through the device for 3 minutes for each wash. Cells were permeabilized by adding 150 μL of 70% ethanol to the inlet and 50 μL of 70% ethanol to the outlet and placing the device for 16 hours at 4°C. After permeabilization, cells were washed three times with PBS and then incubated with 10% formamide in 2X SSC for 30 minutes at 37 °C. Cells were hybridized by adding 50 μL of indicated probes (concentrations discussed below) in buffer containing 2X SSC, 10% formamide, and 100 mg/mL dextran sulfate at 37°c in the dark for 12 hours. Cells were washed twice with 10% formamide in 2X SSC. Finally, cells were counterlabeled with 1 μg/mL Hoechst 33342 and imaged in either PBS or VectaShield mounting media (Vector Labs).

We note here that a key practice to minimize the loss of flow and cells during the smFISH protocol was to implement a simple stepwise procedure that would minimize the loss of hydrostatic pressure in the microfluidic system during fluid replacement. At every buffer exchange step, the outlet was emptied first, then the inlet was emptied and immediately refilled with the new buffer to reestablish flow. All solutions, including the viscous hybridization buffer, passed through the channel and could be displaced within 60 s by addition of a subsequent solution (Fig. S1a in the Supporting Material). We confirmed that the multiple incubation and wash steps did not result in significant cell loss (Fig. S1b).

Labeled cells were imaged on Nikon Eclipse Ti spinning disk confocal microscope with a plan apochromatic 100x oil objective (Nikon; NA 1.45). Regions of interest were re-identified using the fiducial markers. For each field, we acquired z stacks of 30-60 images with 0.3 μm intervals.

Implementing the smFISH protocol in the device, we found that moderate probe concentrations (100 - 200 nM) yielded labeling comparable to that achieved with cells on a dish; higher probe concentrations (500 nM) resulted in high background and the loss of distinct spots (Fig. S1c). In addition, both in conventional culture and in the device, we found very low nonspecific probe binding when performing labeling for transcripts of *GAS6*, a gene not normally expressed in Jurkat cells (16) (200 nM probes; Fig. S1d).

### Image analysis

The Z-center for each cell was manually identified and the mean fluorescence intensity (MFI) for nuclear Ch-RelA in this slice was quantified at every time point. Background was determined by measuring the MFI of an area equivalent to the cell nucleus but in an area of the channel with no objects in the same slice. The background MFI was then subtracted from the nuclear MFI. Cells that were displaced from the traps or divided during imaging were not analyzed. The features for Ch-RelA nuclear translocation were extracted from live-cell time courses as previously reported (7, 12). Fixed cells with labeled transcripts were registered to live cells based on fiducial markers. Cells outlines were traced manually and transcripts were identified and counted using FISH-Quant software (17) as previously described (12).

### Chromatin Immunoprecipitation

EZ ChIP Kit (Upstate) reagents and protocols were used as described previously (12). 15 × 10^6^ cells were fixed in 1% formaldehyde for 10 min. Unreacted formaldehyde was quenched with 125 mM glycine for 5 min. Cells were washed three times with PBS and lysed with 1% SDS lysis buffer in the presence of cOmplete™ protease inhibitor cocktail (Roche). Lysed cells were sonicated with a Diagenode Bioruptor (Settings: 7 cycles of 5 min per cycle with 30 s pulses followed by 30 s of incubation at a high power). Samples were pre-cleared with protein agarose G (Millipore) and 1% of each sample was used as a percent input. Samples were incubated for 16 hours at 4°C. Protein agarose G was then added to the samples and incubated for 1 hour at 4°C. Beads were washed once each with low salt, high salt, and LiCl immune complex wash buffers, and then washed twice with TE (10 mM Tris, 1 mM EDTA, pH 8.0). DNA-protein complexes were eluted from agarose beads with 1% SDS and 100 nM NaHCO3. Crosslinks were reversed by incubating samples with NaCl overnight at 65°C. DNA was purified with DNA spin filters and eluted with TE. DNA was quantified using quantitative PCR (BioRad iCycler, iQ5) using SYBR Green Supermix (BioRad). qPCR was performed in triplicate and melt curves were run to ensure product specificity. Antibodies and primers are listed in table below.

**Table.**
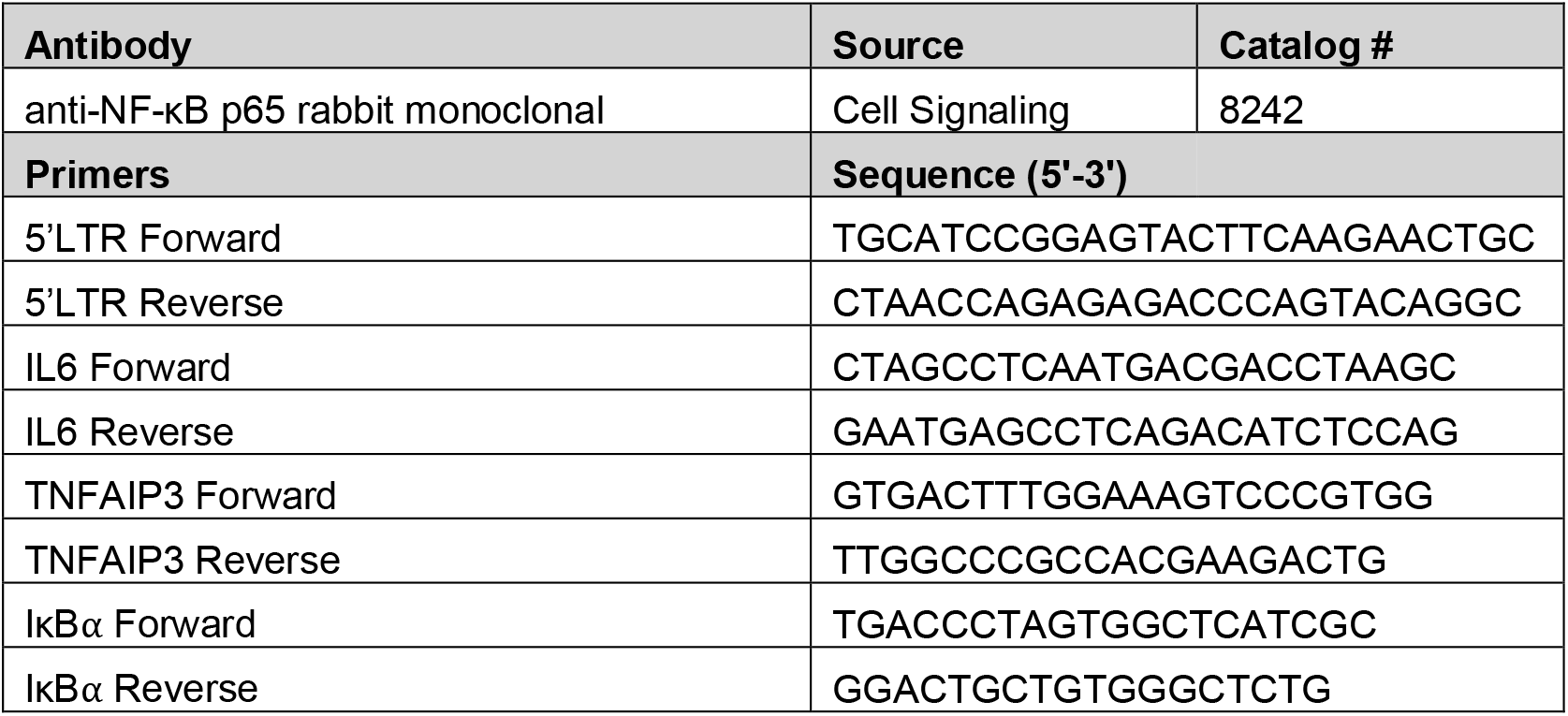
Antibodies and primers

### Computational modeling

To make predictions about fold-change detection, RelA promoter binding, and transcript variance across a range of target transcriptioncal outputs, we used a previously reported computational model of TNF-RelA signaling (7) (Table S1). The transcription rate of the RelA target promoter is given by the following ordinary differential equation:

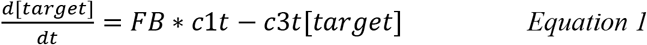

in which the transcript synthesis rate *c1t* is modified by the fractional binding (*FB*) of nuclear RelA (*nRelA*) to the target promoter and the target transcript is degraded at a rate *c3t*. Fractional binding of nRelA depends on the affinity of RelA for the target promoter (*k_RelA_*), the concentration of the competitor protein (*Comp*), the affinity of the competitor for the target promoter (*k_Comp_*), and the Hill coefficient (*h*):

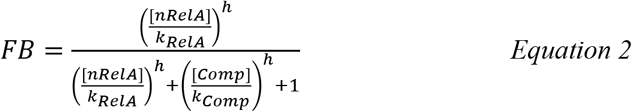

To predict behavior across a range of RelA target promoters with different transcriptional outputs, we first scanned across a range of transcript synthesis and degradation rates sufficient to produce the approximate absolute change in transcription output observed between *NFKBIA* and *IL6*. All other parameters remained the same as previously reported (7). We next varied the Hill coefficient, which effectively varies the steepness of the change in transcriptional response over a rane of nRelA concentration. Finally, we also varied transcriptional output by scanning across a range of competitor affinity sufficient to produce the approximate absolute change in transcription output observed experimentally, as well as a range of competitor affinity for the competitor promoter (to modulate the relative abundance of RelA vs. Comp, which is less well determined). All other parameters, including transcript synthesis and degradation rates, remained the same as previously reported (7).

For each parameter combination, we simulated 450 ‘cells’ by varying total RelA abundance and *k_act_* (a proxy for the strength of activation of pathways leading to IKK), as described previously (7). After setting total RelA abundance, the system is allowed to reach an unstimulated steady state with *k_act_* = 0, setting basal cell conditions, including basal transcript numbers, then stimulation is initiated by switching *k_act_* to its value for this ‘cell’. We then estimated the predicted correlation of number of target transcripts at 120 min vs. maximal fold-change in nRelA over these 120 min. We also estimated the variance of target transcripts at 120 min in the presence of TNF relative to the variance in transcripts in the basal state. Finally, we estimated the average fractional binding on the target promoter for each of combination of competitor affinities at the peak of nRelA concentration at 30 min based on Equation 2.

### Statistical analysis

Differences between transcript distributions were determined by Kolmogorov–Smirnov test (p < 0.05). 95% confidence intervals (CIs) on descriptive statistics of RNA distributions were estimated from the 2.5% and 97.5% quantiles of copy numbers per cell bootstrapped by sampling 1000 times with replacement. For CIs obtained empirically, the difference between two quantities was inferred to be significant if the 95% CIs did not overlap. All statistical tests, regression and correlation analyses were performed in Prism (Graphpad) unless otherwise indicated.

## Results

### Implementation of single molecule RNA FISH for suspension T cells in a passive-flow microfluidic device

We previously developed a user-friendly microfluidic device that immobilizes suspension cells in an array of traps using passive gravity-driven flow between two reservoirs to implement long-term live-cell imaging of suspension cells (13). Our goal here was to collect paired data, tracking NF-κB RelA nuclear translocation dynamics by live-cell imaging and RelA-driven transcriptional output by smFISH in the same cells (Fig. 1a), to gain insight into how NF-κB signaling dynamics are converted to transcriptional outputs in Jurkat T cells. A challenge posed by fixed-cell response assays such as smFISH is that they require multiple fluid exchanges within the device to allow for incubations with different reagents and multiple wash steps. Following optimization of the workflow to implement smFISH in the device (see Methods), imaging Z stacks on the device produced well-defined puncta, representing individual mRNA, which were suitable for downstream analyses (Fig. 1b).

**Fig. 1.**
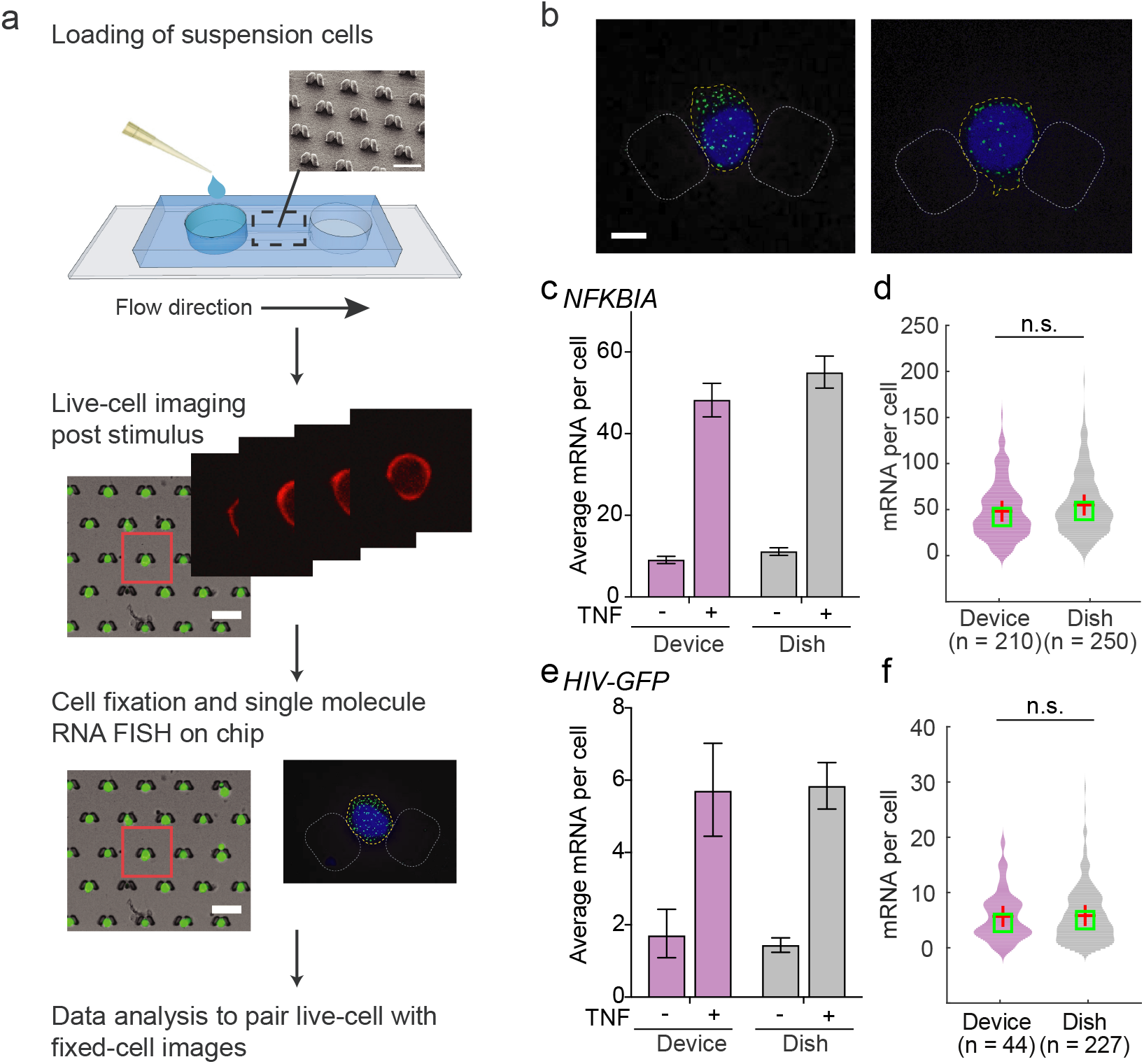
Optimization of a microfluidic device-enabled protocol to connect signaling dynamics to smFISH measurements in single suspension cells. (**a**) Schematic of the live-cell-to-fixed-cell protocol. A cell suspension is pipetted into the inlet of a microfluidic device and flowed into the channel containing traps that catch cells via gravity. Upon stimulation with a media exchange, live cells are imaged in the traps for a desired amount of time. Then cells are fixed and assayed by single-molecule RNA fluorescence in situ hybridization (smFISH). Data is then processed and the live-cell and fixed-cell images are analyzed together for the same single cells. (**b**) Representative images of Jurkat J65c cells treated with 20 ng/ml TNF for 1 hour, fixed in traps and labeled for single *NFKBIA* mRNA with FAM-conjugated probes (green). Nuclei stained with Hoechst (blue); scale bar is 5 μm. (**c**) Bar graph of mean *NFKBIA* transcription for unstimulated cells and cells stimulated with 20 ng/ml TNF for 1 hour measured in the passive-flow device (purple) and in a tissue culture dish (gray). Data are presented as the mean ± 95% confidence intervals obtained by bootstrapping. (**d**) Violin plots of TNF-stimulated *NFKBIA* transcript distributions at 1 hour measured in the passive-flow device (purple) or tissue culture dish (gray). There are no significant differences between the distributions (n.s., p > 0.05 by Kolmogorov-Smirnov, or K.S., test). (**e-f**) Bar graph of mean *HIV-GFP* transcription (e) and violin plots of *HIV-GFP* transcript distributions (f) following 2 hours of TNF stimulation. All other details as in (c-d).

To validate smFISH implementation in the device, we quantified *NFKBIA* transcription in J65c cells, a clonal population of Jurkat T lymphocytes stably expressing an NF-κB mCherry-RelA (Ch-RelA) reporter protein. We compared measurements collected in the device to our previously published measurements of *NFKBIA* transcription collected in a dish (12). We found that mean *NFKBIA* transcript numbers and the distribution of *NFKBIA* transcripts before and one hour after TNF stimulation were indistinguishable between the two formats (Fig. 1c-d). A similar comparison for TNF-stimulated HIV transcripts from clonal J65c Jurkat cell populations containing a latent-but-inducible HIV integration with very low transcript abundance also showed no evidence of changes in transcriptional output between formats (12) (Fig. 1e-f). Overall, our data show that smFISH for both high and low abundance inducible transcripts can be implemented in T cells in our passive-flow device, allowing for a live-cell-to-fixed-cell imaging workflow for non-adherent cells.

### Jurkat cells exhibit fold-change detection of RelA at promoters with high levels of inducible transcription

We proceeded to image TNF-induced NF-κB RelA translocation dynamics followed by smFISH in J65c cells to explore NF-κB RelA signal input–transcriptional output relationships. Ch-RelA translocation was imaged for 2 hours following stimulation with 20 ng/ml TNF (Fig. 2a, left). Cells were then fixed in the traps and transcripts of the RelA-target gene were labeled by smFISH in the device (here for *NFKBIA*) to produce a set of live-cell signaling traces paired with transcriptional output (Fig. 2a, right).

**Fig. 2.**
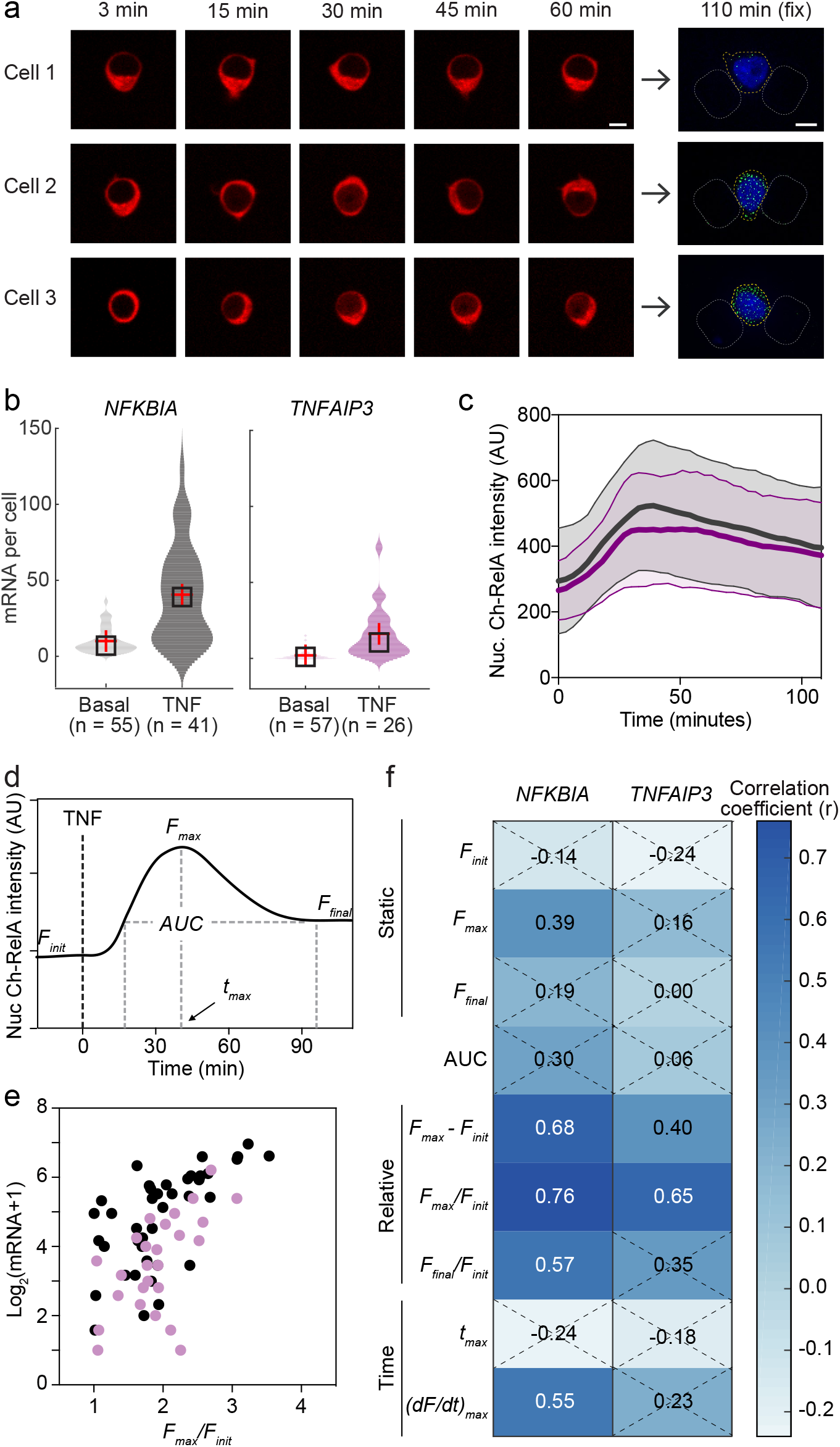
Fold-change detection of nuclear RelA is observed in Jurkat T cells at promoters exhibiting high levels of TNF-inducible transcription. (**a**) Time-lapse images of Ch-RelA paired with smFISH images of *NFKBIA* transcripts at 2 hours after 20 ng/ml TNF treatment from the same cells. Scale bar is 5 μm. (**b**) Violin plots of transcript number distributions for *NFKBIA* (gray) and *TNFAIP3* (purple) before and 2 hours after TNF stimulation in the device. (**c**) Time course traces of nuclear Ch-RelA from J65c cells collected in the passive-flow device after 20 ng/ml TNF treatment for two independent experiments. Data presented as the mean ± SD of individual cell traces (n = 41, traces linked to *NFKBIA*, gray; n = 26, traces linked to *TNFAIP3*, purple). (**d**) Schematic indicating features calculated from individual nuclear Ch-RelA time courses, including initial, maximal, and final fluorescence (*F_i_, F_max_*, and *F_final_*), time *F_max_* is reached (*t_max_*), and area under the curve (*AUC*). (**e**) Scatter plot of transcript number (2 hrs post-TNF addition) versus the maximum fold change in nuclear Ch-RelA (*F_max_*/*F_init_*) for *NFKBIA* (gray) and *TNFAIP3* (purple). (**f**) Heat map of Pearson correlation coefficients with transcript number for all extracted metrics. Non-significant correlations (p > 0.05) indicated by a dashed X.

We first explored input–output relationships for the high abundance targets *NFKBIA* and *TNFAIP3*. Following 2 hours of TNF stimulation, the average number of transcripts increased to 40 and 16 (observed maximum of 124 and 73) transcripts per cell, respectively, for *NFKBIA* and *TNFAIP3* (Fig. 2b). Ch-RelA nuclear translocation over that period was not significantly different between the two experiments (Fig. 2c). We note that TNF-stimulated Ch-RelA translocation dynamics differed somewhat between the device and a tissue culture dish; nuclear Ch-RelA peaked five to ten minutes later and was sustained over a longer period (monitored for 2 hours) in the device versus in a dish (Fig. S2a). This slower and more sustained pattern of nuclear Ch-RelA translocation was not due to a delay in TNF stimulation between the inlet and outlet because the average response of the cells near the inlet of the device was indistinguishable from the response of cells closer to the outlet (Fig. S2b). Despite the slower and more sustained signaling dynamics observed in the device, transcription remained unchanged (Fig. 1c-f), suggesting that T cells respond to a feature of the signaling dynamics that is conserved across formats.

We previously showed that maximum fold change in nuclear RelA is predictive of *NFKBIA* and *TNFAIP3* transcriptional output in HeLa cells (7) and of the fluorescent protein production from an HIV activation reporter construct in Jurkat T cells (12). To evaluate which aspects of RelA signaling are most predictive of *NFKBIA* and *TNFAIP3* transcriptional output in Jurkat cells, we extracted features from individual TNF-stimulated Ch-RelA nuclear intensity time courses (Fig. 2d) and matched them to the number of transcripts produced in the same cell. We included absolute nuclear RelA intensity (initial, maximum, and final), features describing relative changes in nuclear intensity (maximum and final intensity normalized to initial intensity), a feature that integrated nuclear intensity over time (area under the curve, AUC) and a feature that described pathway flux (the maximum rate of change).

When we compared correlations of each Ch-RelA feature with transcript number, we found that the maximum fold change in RelA nuclear intensity (*F_max_*/*F_init_*) was the strongest predictor for *NFKBIA* and *TNFAIP3* transcription levels across all features considered (Fig. 2e-f), consistent with our observations in HeLa cells. Interestingly, we found that the distributions of *F_max_*/*F_init_* for nuclear RelA were the same for translocation time courses collected in the tissue culture dish versus the device (Fig. S2c), which may explain why the observed differences in translocation dynamics between the two formats did not significantly affect transcriptional output. Overall, we conclude that fold-change detection of nuclear RelA in response to TNF stimulation is maintained in Jurkat T cells for the high abundance RelA-target genes *NFKBIA* and *TNFAIP3*.

### Fold change in nuclear RelA is conserved at promoters exhibiting low levels of inducible transcription

We recently studied RelA-mediated activation of latent HIV using J65c cells harboring unique integrations of an HIV lentiviral construct driving expression of GFP and the viral protein Tat. We specifically studied two clones, referred to as J65c 4.4 and 6.6 (12), which do not express virus in the basal state but are activated by TNF. Two hours of TNF treatment with viral positive feedback blocked induced similar, very low mean transcription at these two viral integration sites (Fig. 3a; ~6 transcripts per cell). We also observed a low TNF-induced transcript mean for the cytokine *IL6* (Fig. 3a) consistent with the expectation that *IL6* is not strongly activated by TNF in immune cells (18); however, the increase in response to TNF was nonetheless significant (as demonstrated by non-overlapping 95% CIs; Fig. 3a). Therefore, we wanted to explore if fold-change detection between nuclear RelA and transcript number would be observed at these promoters, which support very low levels of inducible transcription.

**Fig. 3.**
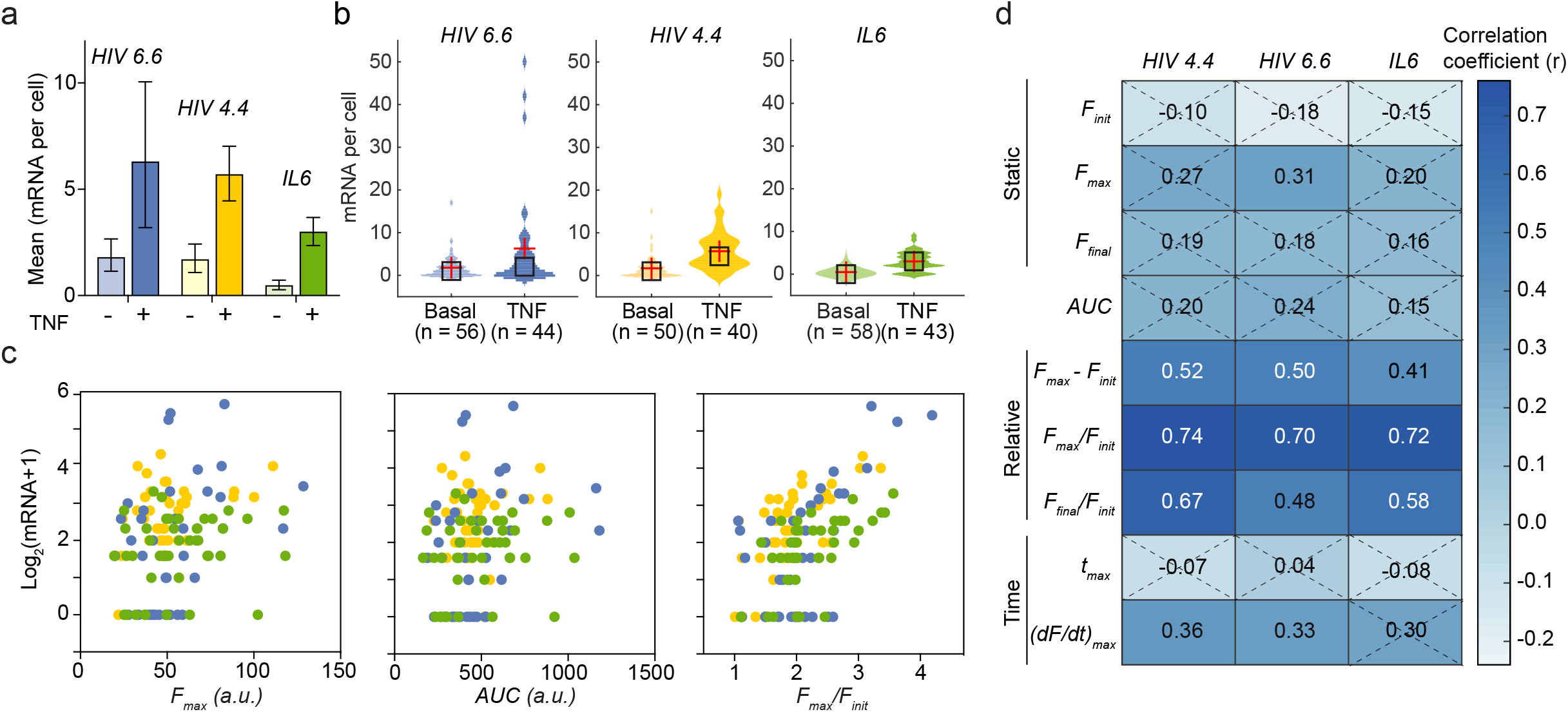
Maximum fold change in nuclear RelA correlates with transcript output for gene promoters exhibiting low levels of TNF-inducible transcription. (**a**) Bar graph of mean transcription of *HIV 6.6* (blue), *HIV 4.4* (yellow), and *IL6* (green) before and 2 hours after 20 ng/ml TNF stimulation. Data are presented as the mean ± 95% CIs obtained by bootstrapping. Significance (*, p < 0.05) inferred by non-overlapping 95% CIs. (**b**) Violin plots of transcript number distributions for *HIV 6.6, HIV 4.4*, and *IL6* before and 2 hours after TNF stimulation. (**c**) Scatter plots of transcript number (2 hrs post-TNF) for *HIV 6.6* (blue), *HIV 4.4* (yellow), and *IL6* (green) versus maximum nuclear F_max_ (left), AUC (middle), and maximum fold change in nuclear Ch-RelA (F_max_/F_init_) with. (**d**) Heat map of Pearson correlation coefficients with transcript number for all extracted metrics. Non-significant correlations (p > 0.05) indicated by a dashed X.

In addition to differences in TNF-induced average transcript numbers, the TNF-induced transcript number distributions also varied across promoters. J65c 6.6 exhibited a highly skewed distribution of HIV transcript numbers in response to TNF stimulation that was more similar to *TNFAIP3*, but J65c 4.4 HIV transcript number distribution and distributions for *IL6* both exhibited much less skew (Fig. 3b and Table 1). We previously observed such differences in transcript distributions and showed that they are associated with differences in the underlying mechanisms of RelA-mediated transcription (12), bringing into question whether the fold-change in nuclear RelA will remain strongly correlated with transcript number at both promoters.

**Table 1.** Statistical moments for experimentally measured transcript distributions. Values for the ± 95% confidence intervals obtained by bootstrapping are listed in parentheses.

| Target | Mean |  | Variance |  | Skewness |  |
| --- | --- | --- | --- | --- | --- | --- |
|  | Basal | + TNF | Basal | + TNF | Basal | + TNF |
| *NFKBIA* | 10.1<br>(8.0, 12.3) | 40.6<br>(31.1, 50.4) | 64.5<br>(36.6, 93.8) | 984.9<br>(596, 1377) | 1.3<br>(0.7, 1.9) | 0.7<br>(0.2, 1.2) |
| *TNFAIP3* | 2.0<br>(1.3, 2.8) | 16.2<br>(10.6, 22.7) | 8.6<br>(2.8, 16.0) | 267.9<br>(78.7, 515.3) | 2.5<br>(1.3, 3.4) | 1.8<br>(0.4, 2.7) |
| *HIV 6.6* | 1.8<br>(1.1, 2.7) | 6.3<br>(3.2, 10.0) | 8.0<br>(1.8, 17.5) | 129.5<br>(17.0, 250.3) | 3.5<br>(0.8, 4.4) | 2.7<br>(1.3, 4.4) |
| *HIV 4.4* | 1.7<br>(1.1, 2.4) | 5.7<br>(4.4, 7.0) | 6.8<br>(1.9, 14.1) | 19.1<br>(10.2, 28.4) | 3.0<br>(1.1, 4.1) | 1.0<br>(0.3, 1.6) |
| *IL6* | 0.5<br>(0.3, 0.7) | 3.0<br>(2.3, 3.7) | 0.8<br>(0.4, 1.3) | 5.0<br>(3.0, 7.1) | 2.0<br>(1.1, 2.8) | 0.5<br>(-0.1, 1.0) |

When we compared correlations of each Ch-RelA feature with transcript number for these low abundance targets, we found that the maximum Ch-RelA nuclear intensity and AUC of Ch-RelA intensity were again only weak predictors of TNF-induced transcript number at 2 hours (Fig. 3c). In contrast, the maximum fold change in RelA nuclear intensity (*F_max_*/*F_init_*) was a strong predictor for *HIV* and *IL6* transcription (Fig. 3c). For *IL6*, the correlation with maximum nuclear RelA fold change held despite the fact that transcript number ranged only from 0 to 10 transcripts per cell. As with high abundance genes, we observed that only features describing relative changes in Ch-RelA nuclear intensity were strong predictors of output, with maximum fold change in nuclear RelA being the strongest predictor of all the features considered (Fig. 3d). Surprisingly, this trend in the correlations was similar for all promoters, despite the differences in their transcript distributions (Fig. 3b and Table 1). Overall, we conclude that RelA fold-change detection is conserved at genes supporting low TNF-inducible mean transcription and with different TNF-induced transcript distributions.

### Mathematical model of I1-FFL RelA signaling recapitulates conservation of fold-change detection at promoters with low inducible transcription

Fold-change detection of signals can be implemented with an I1-FFL signaling motif, one of only two signaling motifs commonly found in biological systems that confers this property (8). We previously implemented an I1-FFL in a mathematical model of TNF-stimulated NF-κB RelA activation and target gene transcription by incorporating a competitor protein that is induced by RelA and competes for binding to promoter κB sites, thereby inhibiting RelA-mediated transcription (7) (Fig. 4a). NF-κB p50, NF-κB p52, and BCL3 are potential candidates for this competitor protein, because they are induced by RelA (19), and they suppress transcription of κB site-containing genes, including *IL6* and the HIV long terminal repeat (20, 21). This model accurately described our observations in HeLa cells for RelA-target genes with high TNF-inducible transcription (7). We now sought to explore if this model could account for our new experimental observations in Jurkat cells extending to RelA-target genes with low TNF-inducible transcription.

**Fig. 4.**
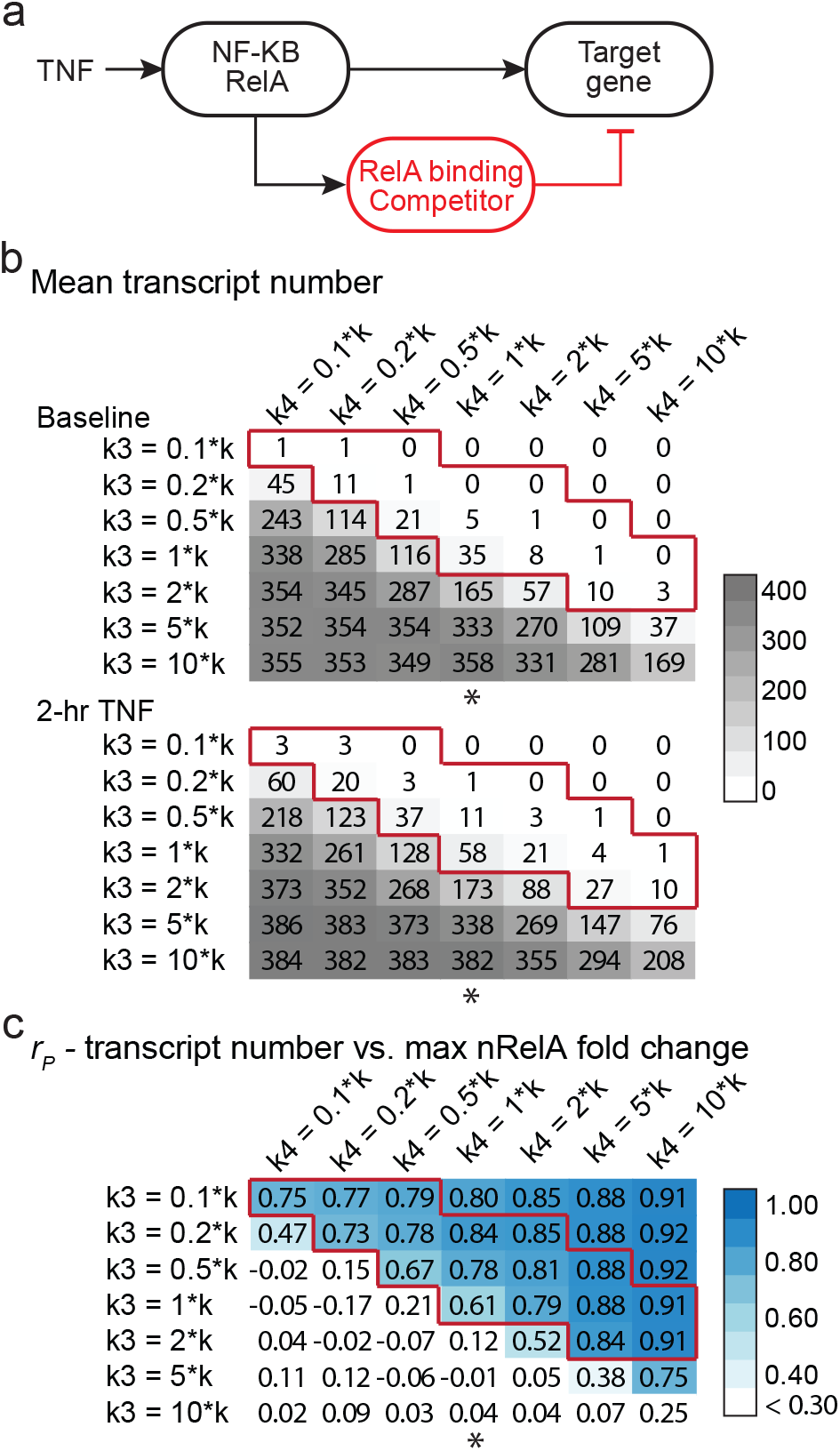
Computational model of NF-κB RelA recapitulates fold-change detection at low abundance transcripts. (**a**) Schematic of how the I1-FFL motif was implemented in a computational model of NF-κB RelA signaling. (**b-c**) Heat maps of the outputs of a parameter scan of competitor affinity (*k3*, columns) and the parameter tuning relative competitor abundance (*k4*, rows) producing a range of (b) transcriptional output before and 2 hours after TNF stimulation (gray) and (c) the corresponding Pearson correlation coefficients of transcript abundance at 2 hours post-TNF with maximum fold-change in nuclear RelA (blue). The ranges that correspond to our experimentally observed absolute change in transcript abundance induced by TNF are outlined in red and the asterisk marks a column discussed in the main text.

We previously showed that a biologically plausible way to tune transcriptional output of RelA targets is to vary the strength of competitor binding affinity relative to RelA binding affinity at its promoter (7), because the relative affinities of RelA and the candidate competitor proteins vary across promoter κB site sequences (22). We performed a model parameter scan to find the range of competitor affinities for target promoters that could reproduce the approximate absolute change in transcript abundance for *NFKBIA, TNFAIP3, HIV* and *IL6* observed in our experiments two hours after addition of 20 ng/mL TNF (Fig. 4b). We also scanned across a range of relative abundances of RelA versus competitor protein levels, which is less well determined and may differ between HeLa and Jurkat cells. All other parameters remained the same as previously reported (7) (Table S1).

We found that moderate to strong correlations between maximum fold-change in nuclear RelA and transcript number were maintained across the entire parameter range considered (Fig. 4c, outlined in red). These simulation results demonstrate that there is a broad parameter range over which the implementation of the I1-FFL in the mathematical model of RelA signaling is consistent with our observation that fold-change detection of nuclear RelA is maintained at promoters with low levels of TNF-inducible transcription.

Target gene transcription in the model could also be affected by transcript synthesis rate, transcript degradation rate, and a Hill coefficient determining the steepness of transcription response. In fact, previous studies showed that specificity in dynamics of transcript abundance for genes responding to the same transcription factor can be achieved by varying the transcript degradation rate (23–25). We therefore performed another parameter scan across a range of transcript synthesis and degradation rates and Hill coefficients and examined the distributions of transcript abundance outputted by the model. Within this scan, it was difficult to find a range of values that reproduced the approximate absolute change in transcript abundance observed experimentally two hours after addition of 20 ng/mL TNF, for the low abundance targets *HIV* and *IL6* (Fig. S3a). Nevertheless, within this limited range, the fold-change correlation was recapitulated (Fig. S3b). Based on these simulation results, we conclude that varying transcriptional output by varying relative competitor affinity and abundance, and to a lesser extent by varying transcript synthesis and degradation rates or the Hill coefficient, can recapitulate fold-change detection at low abundance targets.

### Model predictions and experiments suggest that changes in competitor alone are not sufficient to explain low vs. high abundance targets driven by fold-change detection

After confirming that the model can accurately recapitulate fold-change detection across a wide range of transcript levels by varying competitor affinity and abundance, we wanted to test additional model predictions for low abundance targets to explore the robustness of the underlying mechanistic assumptions in the model. Specifically, cell-to-cell heterogeneity in the model is produced by varying the initial concentrations of total RelA and of TNF-induced IKK activity across cell simulations, which generates a distribution of transcript numbers because transcription is proportional to RelA binding at the target promoter (7). Therefore, we next explored how the predicted transcript distributions compared to our experimental measurements across a range of transcriptional outputs.

Scanning across the same parameter range explored in Fig. 4b-c, we found that the model predicted that the variance in transcript number following TNF treatment (normalized to the variance in unstimulated conditions) generally increased as TNF-induced target transcript abundance decreased (Fig. S4a). When we chose a single set of parameters that produced average TNF-stimulated mRNA abundance ranging from 58 to 1 transcript per cell (Fig. 5a, top), which encompasses the range of transcription observed for the five targets included in this study (Fig. 5b, top), a monotonically increasing trend was clear (Fig. 5a, center). In contrast, although we observed an increase in relative variance between *NFKBIA* and *TNFAIP3*, we observed that the relative variances of TNF-induced *HIV* and *IL6* decreased as compared *TNFAIP3* (Fig. 5b, center). Varying transcriptional output by varying synthesis and degradation rates also failed to reproduce the observed trend in relative variance (Fig. S4b). Thus, the model does not accurately reproduce trends in TNF-induced transcript distributions, even though it accurately predicted fold-change detection of RelA for promoters across a range of transcriptional outputs achieved by varying relative competitor affinities (Fig. 4c).

**Fig. 5.**
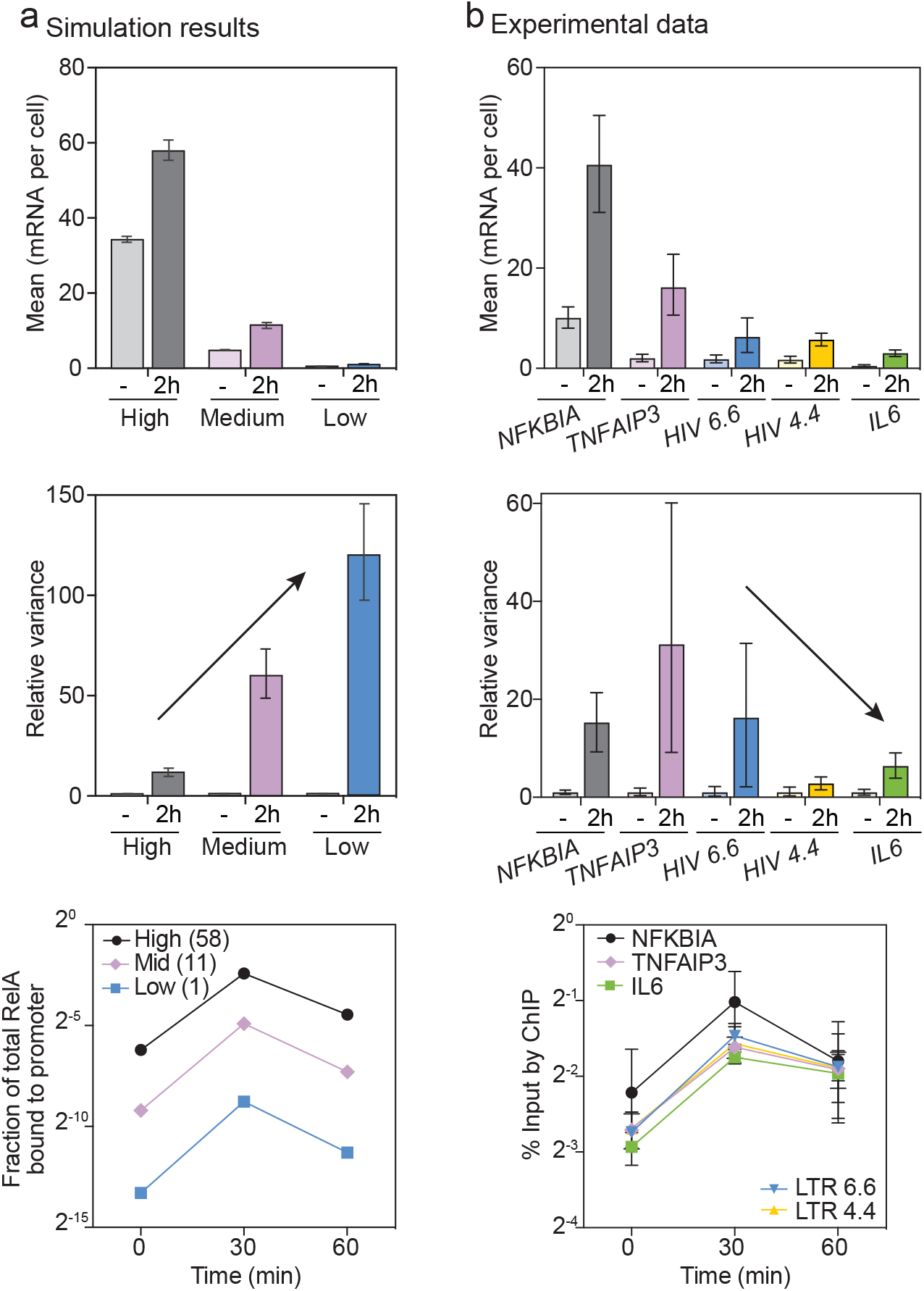
Tuning transcriptional output across promoters by increasing competitor binding does not match experimentally observed trends in transcriptional noise and RelA binding. (**a**) Simulation results, showing a bar graph of mean transcriptional output before and 2 hours after TNF stimulation (top), a bar graph of relative variance calculated as the variance of transcript number normalized to the variance of transcript number in the unstimulated state (center), and time courses of mean fraction of total RelA bound to promoter (bottom). Simulations were run for for k4 = 1*k and k3 = 1*k (high abundance, gray), 0.5*k (medium abundance, purple), and 0.2*k (low abundance, blue). Error bars represent the ± 95% CIs obtained by bootstrapping (error bars are smaller than the markers in the bottom graph). (**b**) Experimental data, showing bar graphs of mean mRNA per cell (top) and relative variance in transcript number (normalized to variance measured in unstimulated cells, center), for cells before or after 2 hours of 20 ng/ml TNF stimulation. Transcript number was measured by smFISH for the 5 indicated targets and error bars indicate ± 95% CIs obtained by bootstrapping. Time courses of enrichment of RelA (% input measured by ChIP) at the 5 target promoters after TNF addition. We show mean ± s.e.m. of independent biological duplicate (LTRs) or triplicate (endogenous promoters); no significant differences in time course as calculated by two-way ANOVA.

Decreasing transcriptional output by increasing competitor affinity relative to RelA affinity (moving upward within a column, Fig 4e**) results in the related prediction that the fraction of RelA bound also decreases (Fig. S5a), which can be tested experimentally. For example, to reduce average TNF-stimulated mRNA levels from 58 to 1 (the same range explored above), the model predicted that the fraction of RelA bound at 30 minutes after TNF treatment would be reduced nearly 80-fold (Fig. 5a, bottom). We used chromatin immunoprecipitation (ChIP) to measure RelA binding at the promoters at 0, 30, and 60 minutes after TNF treatment. In contrast to the model prediction, we found no significant differences in RelA binding across all promoters following TNF treatment (Fig. 5b, bottom). Based on these results, we conclude that competitor affinity alone cannot explain the difference in transcript abundance at low vs. high output targets.

Overall, our quantitative single-cell data set of NF-κB signal-response relationships for low abundance transcripts provided additional constraints for testing TNF-RelA model predictions. We conclude that to maintain fold-change detection across promoters with widely varying transcriptional outputs requires changes in multiple determinants of transcriptional outputs or additional mechanisms to modulate transcript abundance.

## Discussion

Signal–response relationships measured in single cells provide insights into regulatory mechanisms. We previously discovered that RelA-target promoters exhibit fold-change detection of TNF-stimulated signaling; here we extended this finding to show that fold-change detection is conserved across cell types and even at promoters with low TNF-induced transcription. Acquiring signal-response relationships in single Jurkat T cells required development of a microfluidic device-enabled imaging protocol to measure live-cell signaling dynamics and fixed-cell transcriptional output in the same suspension cell.

Our findings suggest that fold-change detection (FCD) may be a general mechanism of TNF-induced NF-κB signal detection. FCD has also been observed in the extracellular regulated kinase pathway (26) and for cyclic adenosine 3’,5’-monophosphate (cAMP) in social amoebae (27), indicating that FCD may confer broadly useful biological functionality. For example, FCD has been proposed as a mechanism by which cells buffer against stochastic variation in signaling molecules by allowing transcriptional output to correlate with signal strength instead of absolute abundance of the signal-driving protein (7, 8). Our results show that this buffering is observed for low abundance transcripts, even though gene expression noise has been observed to increase as mean expression decreases (28, 29).

Our results also demonstrate the advantages of measuring transcript number, rather than a transcriptional reporter, when inferring signal-response relationships from single-cell data. We previously quantified how TNF-stimulated RelA signaling features correlated to levels of HIV activation in J65c 4.4 and 6.6 using a GFP gene expression reporter that is subject to positive feedback amplification by the HIV encoded protein Tat (12). A major weakness of that approach was that only a minor fraction of the J65c 4.4 cells (25%) exhibited detectable gene expression because transcription from this promoter was not efficiently amplified by viral positive feedback (12). While we did find that the maximum fold change in RelA was the strongest predictor of the Tat-mediated GFP protein expression for both viral integrations, we observed that this correlation was significantly weaker for J65c 4.4 versus J65c 6.6 (r = 0.55 versus r = 0.78). By directly measuring transcriptional output using smFISH, we now conclude that the RelA signal-to-transcript relationship is identical for both clones, and inefficient Tat-mediated positive feedback degrades the correlation for J65c 4.4.

To gain mechanistic insight, we compared our experimental measurements across a range of transcript abundances to simulations of a mathematical model of TNF-mediated NF-κB signaling (7). By scanning across a range of values for parameters affecting the abundance of the target transcript, we found that no single parameter could successfully vary transcriptional output while simultaneously reproducing all the trends in our experimental data. Tuning relative competitor affinity recapitulated our observation of fold-change detection across cells for low abundance transcripts (Fig. 4), but it did not recapitulate transcriptional noise trends at the low abundance targets (Fig. 5). This finding was in contrast to our previous findings made for targets with higher transcript abundance (7), highlighting the additional information gained by exploring behavior at low abundance targets.

Why might there be limited transcription at some promoters, despite similar transcription factor occupancy and transcript stability? One potential explanation is differences in the local chromatin environment. Genome-wide studies of transcriptional reporters integrated at different genomic locations, including the HIV LTR, show that transcriptional activity is affected by features of the local chromatin environment (30, 31) and the stability of nucleosomes, which compact DNA and may inhibit access of transcriptional machinery (32). TF-induced chromatin and nucleosome remodeling appears to be a rate-limiting step for transcriptional activation (33, 34). Because rates of chromatin and nucleosome remodeling are slower than TF binding, the chromatin state may decode TF activity on a time scale that is independent of occupancy (35). Such behavior cannot be captured in a mechanistic model by simply tuning a synthesis rate by fractional TF binding.

Chromatin remodeling has been observed to be rate limiting for RelA-mediated transcription (36). Although RelA cannot directly remodel chromatin, RelA recruits the histone acetyltransferase (HAT) p300, which acetylates histones to enhance transcriptional activity (21, 37). Moreover, candidate competitor proteins p50, p52, and BCL3, mediate their repressive transcriptional activity at the HIV LTR and other RelA-target promoters by recruiting histone deacetylases (38–40). Thus, RelA has the potential to remodel chromatin and thus change transcriptional output on a time scale that is different than the lifetime of the RelA-bound promoter complex.

Going forward, quantitative models of transcription activated by the NF-κB pathway, or other gene regulatory pathways, will need to incorporate many contributing factors (i.e., signal decoding, promoter architecture, and chromatin environment) to accurately predict input-output relationships for specific genes in individual cells of different types. The need to describe the recruitment of cofactor complexes and changes in chromatin environment at the promoter has been addressed mathematically by modeling promoters as switching between discrete states (41–45). A recent study demonstrated how a transcription factor that affects the frequency of transitions between promoter states can cause strong transcriptional responses even with weak binding (46). We recently showed that TNF can affect the frequency of transitions between promoter states or the size of the transcriptional burst in the active promoter conformation, depending on the chromatin state of the promoter (12). Accurately modeling how transcription factors influence switching between these states will require additional direct observations of signaling input-output relationships across cell types, signaling pathways, and other gene targets, as well as coupling with measurements and manipulations of additional regulatory factors such as chromatin environment and transcriptional co-factor binding.

## Acknowledgements

This work was funded by the National Science Foundation (CBET-1454301 to KMJ) and the NIH (R01-GM104247 to SG). V.C.W. was supported by an NIH Predoctoral Training Grant in Genetics (2T32GM007499-36, 5T32GM007499-34, 5T32GM007499-35).

